# Prior precision modulates the minimisation of prediction error in human auditory cortex

**DOI:** 10.1101/415083

**Authors:** Yi-Fang Hsu, Florian Waszak, Jarmo A. Hämäläinen

**Affiliations:** Department of Educational Psychology and Counselling, National Taiwan Normal University, 10610 Taipei, Taiwan; Institute for Research Excellence in Learning Sciences, National Taiwan Normal University, 10610 Taipei, Taiwan; Université Paris Descartes, Sorbonne Paris Cité, 75006 Paris, France; CNRS (Laboratoire Psychologie de la Perception, UMR 8242), 75006 Paris, France; Jyväskylä Centre for Interdisciplinary Brain Research, Department of Psychology, University of Jyväskylä, 40014 Jyväskylä, Finland

**Keywords:** predictive coding, prediction error, auditory perception, repetition, magnetoencephalography (MEG).

## Abstract

The predictive coding model of perception proposes that successful representation of the perceptual world depends upon cancelling out the discrepancy between prediction and sensory input (i.e., prediction error). Recent studies further suggest a distinction between prediction error associated with non-predicted stimuli of different prior precision (i.e., inverse variance). However, it is not fully understood how prediction error from different precision levels is minimised in the predictive process. The current research used magnetoencephalography (MEG) to examine whether prior precision modulates the cortical dynamics of the making of perceptual inferences. We presented participants with cycles of repeated tone quartets which consisted of three prime tones and one probe tone. Within each cycle, the three prime tones remained identical while the probe tones changed at some random point (e.g., from repetition of 123X to repetition of 123Y). Therefore, the repetition of probe tones can reveal the development of perceptual inferences in low and high precision contexts depending on its position within the cycle. We found that the two conditions resemble each other in terms of N1m modulation (as both were associated with N1m suppression) but differ in terms of N2m modulation. While repeated probe tones in low precision context did not exhibit any modulatory effect, repeated probe tones in high precision context elicited a suppression and rebound of the N2m source power. The differentiation suggested that the minimisation of prediction error in low and high precision contexts likely involves distinct mechanisms.

## 1.1 Introduction

Our brain constantly predicts forthcoming sensory inputs. The predictive coding model of perception postulates that perception entails two distinct neurocomputational components, the top-down propagation of prediction and the bottom-up propagation of prediction error (Rao & Ballard, 1999; Friston, 2005, 2009; Summerfield et al., 2008; see Clark, 2013 for a review). The flow of information takes place between multiple hierarchical levels harbouring both representational units and error units (Egner et al., 2010). While the representational units encode prediction about the causal structure of the environment and feed it backward to the next lower level, the error units encode the discrepancy between prediction and sensory input as prediction error and communicate it forward to the next higher level. The message-passing between hierarchical cortical levels iterates to match prediction and sensory input as much as possible, that is, to minimise prediction error in the system.

Recent research further suggested the necessity to distinguish between two conditions inducing prediction error: the unpredicted condition (where there is no precise prediction) and mispredicted condition (where there is a precise prediction being violated). Conceptually, unpredicted condition is mainly associated with prediction error generated by sensory input that is not anticipated, whereas mispredicted condition triggers not only prediction error generated by sensory input that is not anticipated but also prediction error generated by prediction that is not fulfilled (Arnal & Giraud, 2012). The dissociation was supported by electroencephalography (EEG) evidence demonstrating that unpredicted and mispredicted stimuli are associated with different amounts of cortical activity (Hsu et al., 2015, 2018). Relative to predicted stimuli, unpredicted stimuli are associated with smaller neuronal response whereas mispredicted stimuli are associated with larger neuronal response on the N1 event-related potential (ERP) component, which is typically considered an electrophysiological indicator for automatic predictive processing (see Bendixen et al., 2012 for a review).

The result pattern can be interpreted in terms of how prediction error is adjusted depending on the precision of the sensory input (Friston, 2005, 2009). Precision refers to the inverse of a signal’s variance, which quantifies the degree of certainty about the signals in general statistical usage (Feldman & Friston, 2010; Ransom et al., 2017). Unpredicted stimuli, relative to mispredicted stimuli, are embedded in contexts of larger variance (i.e., lower precision); therefore, prediction error is weighted less in the former than the latter (Schröger et al., 2015). Such precision weighting mechanism is suggested to be encoded by gain in superficial pyramidal cells in sensory cortices (Feldman & Friston, 2010; Fardo et al., 2017). The idea also conforms to previous research on the mismatch negativity (MMN) which reported a significant difference when contrasting between a deviant sound embedded in an equiprobable sequence (i.e., a low precision context) and a deviant sound embedded in a standard sequence (i.e., a high precision context) (e.g., Jacobsen & Schröger, 2001; see Näätänen et al., 2005 for a review; but see Ahmed et al., 2011; Astikainen et al., 2011; Nakamura et al., 2011 versus Farley et al., 2010; Fishman & Steinschneider, 2012; Kaliukhovich & Vogels, 2014 for an ongoing debate in animal research).

The differentiation raised the question whether the minimisation of prediction error in low and high precision contexts also involves distinct mechanisms. Here we looked into the neuronal underpinnings of the minimisation of prediction error in low and high precision contexts using magnetoencephalography (MEG). Specifically, we examined whether there is a difference between the two conditions in terms of N1m and N2m modulation, given that these long-latency components are mediated by top-down effects in cortical networks and therefore rest on backward connections (Garrido et al., 2007). We presented participants with cycles of repeated tone quartet which consisted of three prime tones and one probe tone. Within each cycle, the three prime tones remained identical while the probe tones changed at some random point (e.g., from repetition of 123X to repetition of 123Y). Therefore, the repetition of probe tones can reveal the development of perceptual inferences in low and high precision contexts depending on its position within the cycle. In the beginning of a cycle where the three prime tones are of little predictive value, the presentation of probe tone X triggers prediction error in a low precision context (because listeners would predict a probe tone to be presented but cannot be quite sure of its frequency). In the middle of a cycle where the prime tones are already associated with probe tone X, the presentation of probe tone Y triggers prediction error in a high precision context (because listeners would tend to predict probe tone X to appear but such expectation is violated). If the minimisation of prediction error in low and high precision contexts involves distinct mechanisms, the repetition of tone quartets 123X and 123Y should modulate MEG responses in different manners.

## 1.2. Materials and methods

### 1.2.1. Participants

Eighteen healthy adults (average age: 24; 6 males; 14 right-handed) with no history of neurological, psychiatric, or visual/hearing impairments as indicated by self-report participated in the experiment. Participants gave written informed consent and were paid for participation. Ethical approval was granted by the ethics committee of National Taiwan Normal University (Taiwan) and the University of Jyväskylä (Finland). Four participants were excluded from data analysis for excessive measurement noise, leaving fourteen participants in the final sample (average age 24; 3 males; 12 right-handed).

### 1.2.2. Stimuli

Sinusoidal tones with a loudness of 80 phons (i.e. 80 dB for tones of 1000 Hz) were generated using Matlab. The duration of each tone was 50 ms (including 5 ms rise/fall times). The frequency of each tone was within the range of 261.626 - 493.883 Hz, matching the absolute frequency of a series of seven natural keys on a modern piano (i.e., C4 D4 E4 F4 G4 A4 B4).

A total of 90 pairs of tone quartets (consisting of three prime tones and one probe tone) were created. Each pair of tone quartet was identical in the prime tones but different in the probe tone in terms of frequency (e.g., F4-E4-G4-**A4** and F4-E4-G4-**D4**). The frequency of the prime tones was determined by a random sampling without replacement, with the exception of any continuously rising or falling sequence to avoid the step inertia expectation (i.e., the expectation that the frequencies of upcoming tones continue in the same direction when the frequencies of previous tones are presented as a scale; Lange, 2009). The frequency of the probe tone can be anything except that of the prime tones.

### 1.2.3. Procedures

A total of 10 blocks of 9 cycles were presented. Each cycle consisted of the repetition of a pair of tone quartet, where the first tone quartet was repeated 4 to 6 times before the second tone quartet was repeated 4 to 6 times. The reason we presented each tone quartet 4 to 6 times was to prevent participants from learning high-order regularities (e.g., correctly anticipating a change in probe tone). Therefore, a cycle could contain 8 to 12 tone quartets. While the repetition of the first tone quartet turned the initially non-predicted probe tone into a predicted tone in a low precision context, the repetition of the second tone quartet turned the initially non-predicted probe tone into a predicted tone in a high precision context (**Figure 1A**).

**Figure 1.**
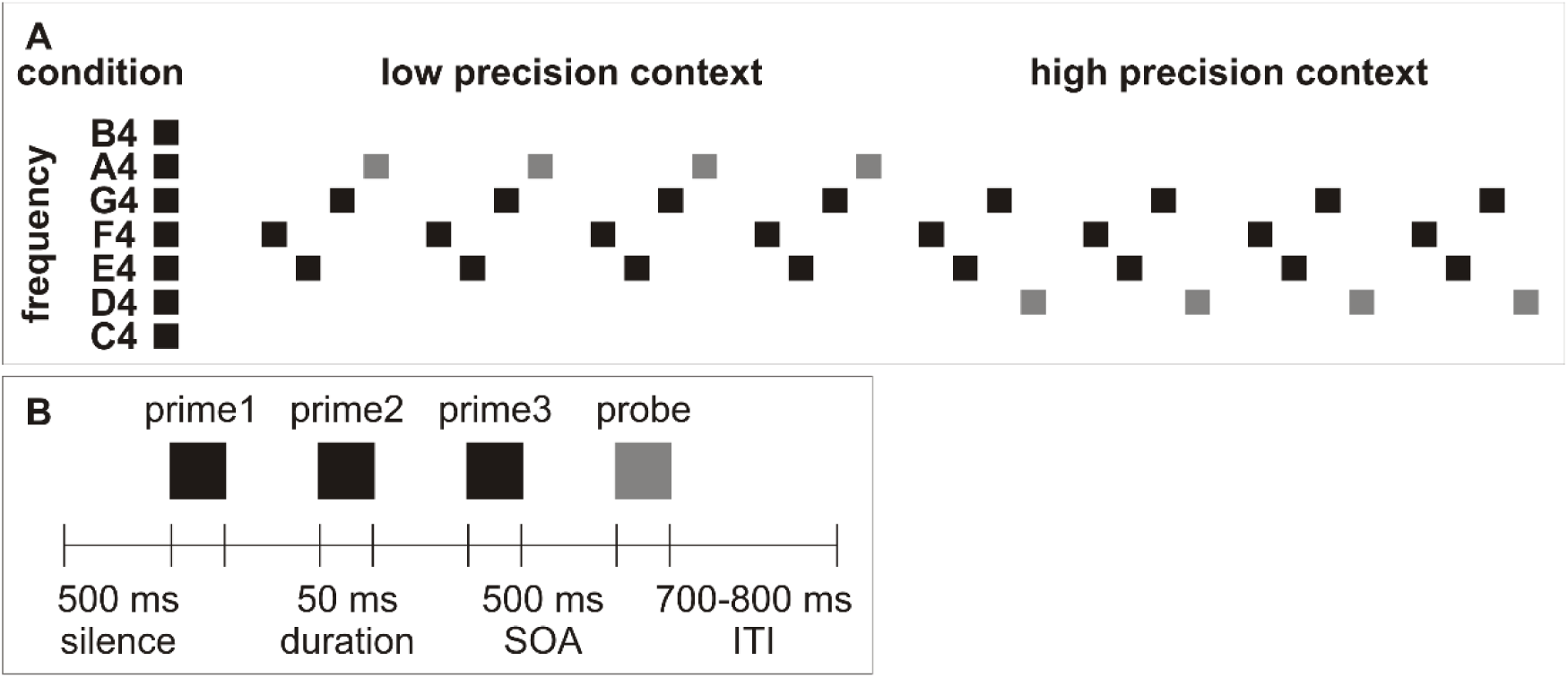
**A.** Schematic representation of a cycle. In this example, the first tone quartet (F4-E4-G4-A4) was repeated 4 times before the second tone quartet (F4-E4-G4-D4) was repeated 4 times. **B.** Schematic representation of a tone quartet.

**Figure 1B** illustrates a tone quartet, which started with a silent interval of 500 ms. Each tone was separated by a 500 ms stimulus onset asynchrony (SOA). 10% of the probe tones were of attenuated loudness by 20 dB. Participants were required to press a key when they detected a softer probe tone as soon as possible to maintain their attention. The offset of the probe tone was followed by a jittered inter-trial interval (ITI) of 700-800 ms. There was no separation between cycles distinct from the ITI. A fixation cross remained on the screen for the duration of the block. The whole experiment took around 42 minutes (i.e., 900 trials x 2800 ms). Presentation (Neurobehavioral Systems, Inc., USA) was used for stimulus presentation. Stimulation was randomised individually for each participant and delivered through two panel speakers situated to the left and right of the participant.

### 1.2.4. Data recording and analysis

MEG data was collected using a 306 channel whole-head device (Elekta Neuromag, Finland) in a two-layered magnetically shielded room at the University of Jyväskylä. The sampling rate was 1000 Hz. A high-pass filter of 0.03 Hz and a low-pass filter of 200 Hz were used. Continuous head position monitoring was used based on five Head-Position Indicator (HPI) coils, with three at the forehead and two behind the ears. Electrooculography (EOG) was recorded using electrodes lateral to each eye and above and below the left eye.

Offline, head movements were corrected and external noise sources were attenuated using the temporal extension of the source subspace separation algorithm (Taulu et al., 2005) in the MaxFilter program (Elekta Neuromag, Finland).

After the initial head movement correction, the data was analysed using BrainStorm 3.2 (Tadel et al., 2011). Signal subspace projection was used to correct for eye blinks. The MEG signal was filtered at 1-40 Hz and segmented from -100 ms to 500 ms relative to the onset of the stimulus using a 100 ms pre-stimulus baseline. Segments with over 5000 fT/cm peak-to-peak values in gradiometers or 7000 fT peak-to-peak values in magnetometers were rejected. As all tone quartets were repeated at least 4 times, segments to the 5th and 6th presentations of tone quartets were also rejected to ensure our analysis is based on equal number of trials. The trial numbers after artefact rejection in each condition are listed in **Table 1**.

**Table 1.** Range, mean, and standard deviation (SD) of trial numbers after artefact rejection in each condition.

| Presentation |  | Low precision context |  |  |  | High precision context |  |  |  |
| --- | --- | --- | --- | --- | --- | --- | --- | --- | --- |
|  | Tone | Min | Max | Mean | SD | Min | Max | Mean | SD |
| 1st | Prime1 | 67 | 90 | 84.43 | 6.89 | 67 | 90 | 84.71 | 6.65 |
|  | Prime2 | 66 | 90 | 83.71 | 7.60 | 64 | 90 | 84.14 | 7.47 |
|  | Prime3 | 59 | 90 | 83.79 | 8.74 | 64 | 90 | 82.93 | 8.32 |
|  | **Probe** | **55** | **85** | **75.57** | **8.28** | **56** | **83** | **75.50** | **7.29** |
| 2nd | Prime1 | 67 | 90 | 83.79 | 7.28 | 62 | 90 | 82.86 | 8.14 |
|  | Prime2 | 68 | 90 | 84.50 | 6.93 | 63 | 90 | 84.50 | 7.43 |
|  | Prime3 | 64 | 90 | 83.79 | 8.14 | 64 | 90 | 84.43 | 7.58 |
|  | **Probe** | **65** | **83** | **76.71** | **5.46** | **59** | **84** | **76.50** | **7.10** |
| 3rd | Prime1 | 67 | 90 | 83.50 | 6.79 | 67 | 90 | 84.36 | 5.85 |
|  | Prime2 | 71 | 90 | 83.86 | 6.19 | 61 | 90 | 83.21 | 8.14 |
|  | Prime3 | 69 | 90 | 83.50 | 7.01 | 66 | 90 | 84.29 | 6.84 |
|  | **Probe** | **58** | **83** | **74.29** | **6.62** | **59** | **82** | **75.93** | **6.22** |
| 4th | Prime1 | 69 | 90 | 84.36 | 6.52 | 66 | 90 | 84.36 | 6.28 |
|  | Prime2 | 71 | 90 | 84.50 | 6.27 | 70 | 90 | 84.64 | 6.18 |
|  | Prime3 | 72 | 90 | 84.36 | 6.21 | 62 | 90 | 82.86 | 8.39 |
|  | **Probe** | **64** | **84** | **76.00** | **6.61** | **58** | **81** | **76.57** | **6.43** |

The experimental effects were examined in source space. As individual magnetic resonance images (MRI) were not available from the participants, the ICBM152 MRI template was used. The weighted minimum norm estimates (wMNE) were calculated using the unconstrained option to allow free orientation of the dipoles in relation to the cortical surface. Three shell spherical head model was used. The wMNE solution was restricted to the cortex. Noise covariance matrix was calculated from the baseline of the averaged responses.

To extract the N1m and N2m measures, we first identified the N1m and N2m from the grand average global field power (GFP) of the gradiometers (across 14 participants and 32 conditions) (**Figure 2A**). Then, we identified brain regions from the Dessikan-Killiany parcellation which showed the largest source activity around the auditory cortices at the N1m and N2m, including transverse temporal, superior temporal, middle temporal, supramarginal, and postcentral regions (**Figure 2B**).

**Figure 2.**
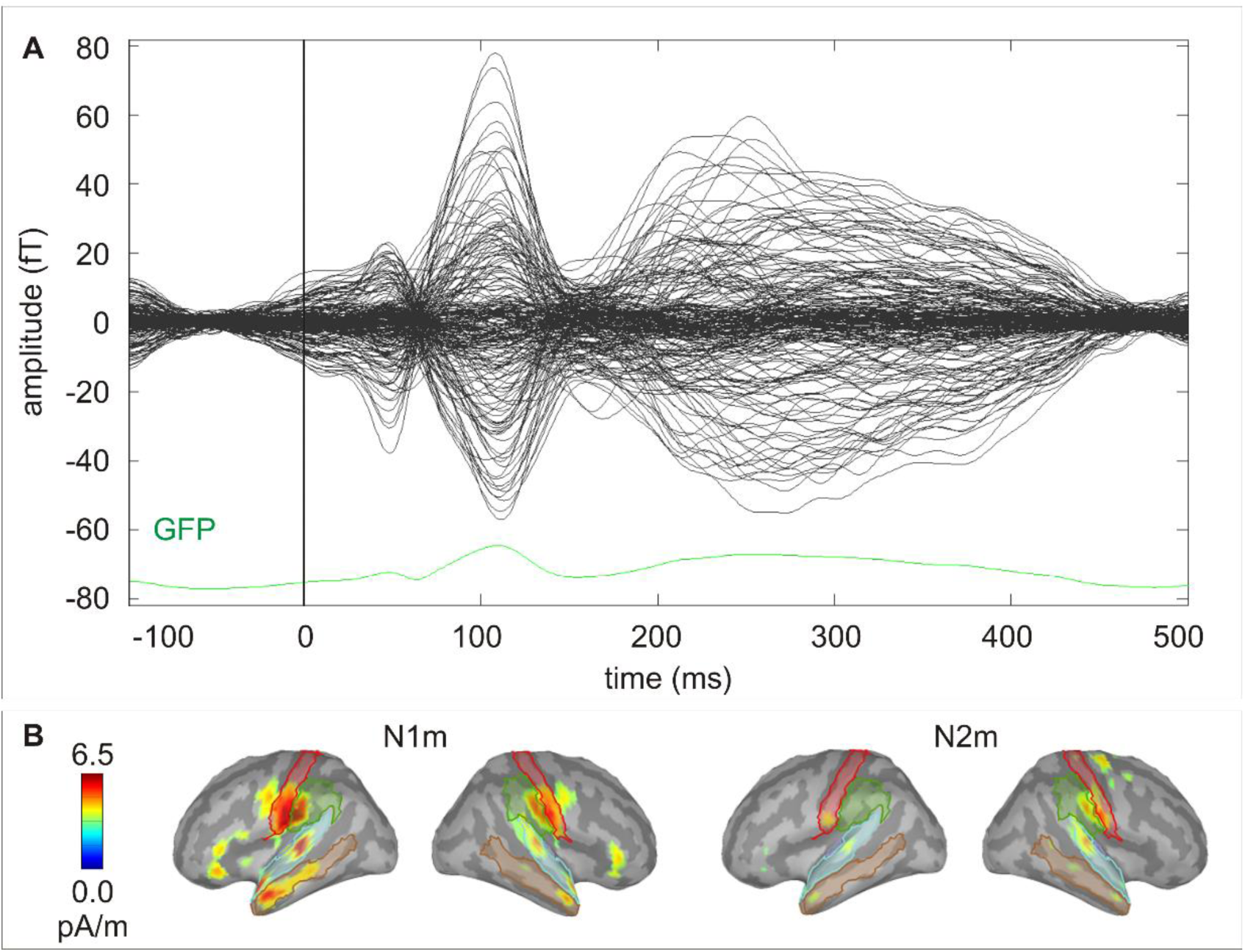
**A.** Butterfly plot of the grand average signals of the gradiometers and the grand average GFP of the gradiometers (across 14 participants and 32 conditions). **B.** Grand average source activity (across 14 participants and 32 conditions) at the N1m and N2m peak. The coloured lines mark the outlines of the 5 brain regions selected to represent the auditory response.

The grand average source solution (across 2 hemispheres, 5 brain regions, 14 participants, and 32 conditions) was used to identify the N1m and N2m time windows for statistical analysis. N1m peak was at ca. 110 ms (time window 85 - 135 ms) and N2m peak was at ca. 220 ms (time window 170 - 270 ms) (**Figure 3**).

**Figure 3.**
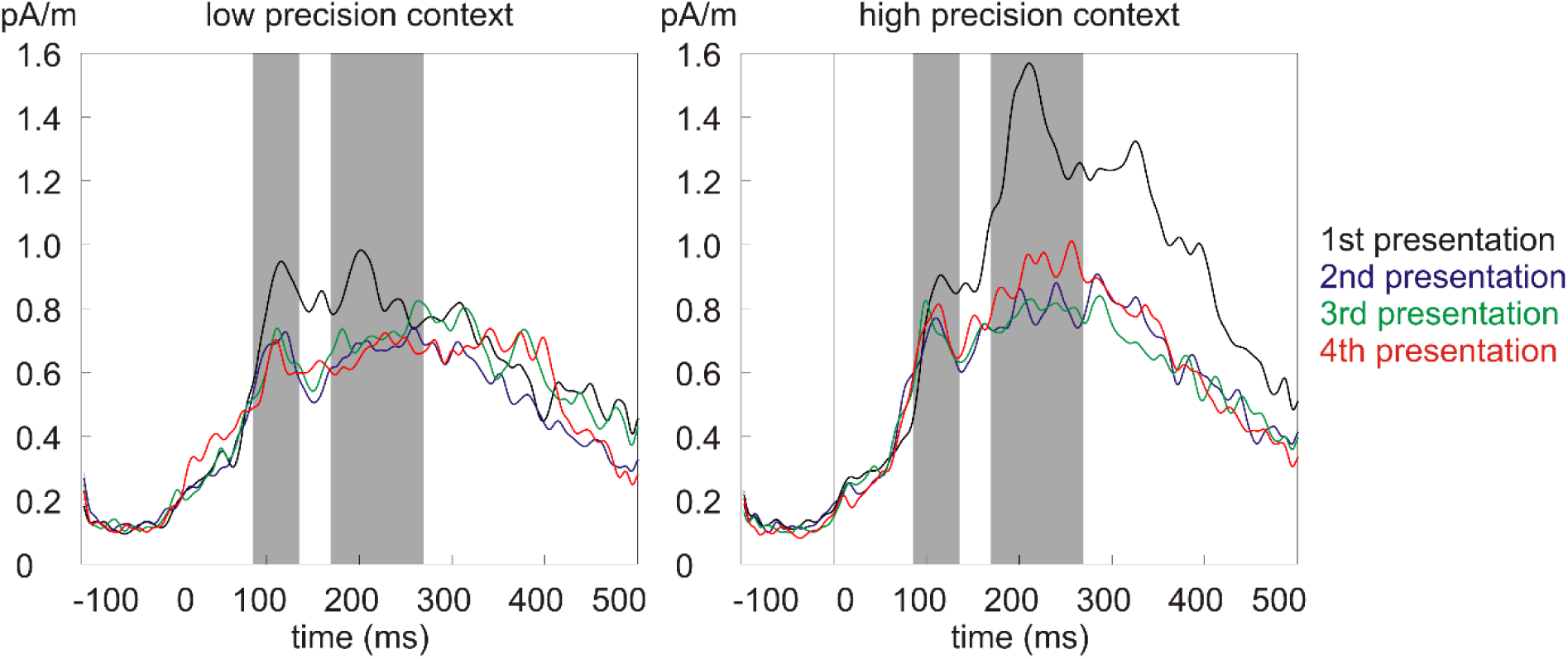
The grand average source waveforms for the four presentations in the low (left) and high (right) precision contexts. Grey bars indicate the N1m and N2m time windows for statistical analysis.

The source power in the N1m and N2m time windows of the probe tones were submitted to the 2 (precision: low/high precision context) x 4 (repetition: 1st/2nd/3rd/4th presentation) repeated measures analysis of variance (ANOVA). Greenhouse-Geisser correction was applied when appropriate (and will be indicated in the following section with epsilon values).

## 1.3. Results

The ANOVA on the N1m source power showed only a main effect of repetition (F(3,39) = 6.33, p < 0.01, partial eta squared = 0.33) (**Figure 4** left). The effect was due to larger response to the 1st presentation compared to all the other presentations (**Table 2**). No significant differences were found between the response strength for the other presentations.

**Table 2.** Results of post hoc paired samples t-tests looking into the main effect of repetition on N1m source power.

| Pair | Mean | SD | 95% Confidence Interval of the Difference |  | t | df | Sig. (2-tailed) |
| --- | --- | --- | --- | --- | --- | --- | --- |
|  |  |  | Lower | Upper |  |  |  |
| 1-2 | 0.13 | 0.18 | 0.03 | 0.24 | 2.68 | 13 | 0.019 |
| 1-3 | 0.14 | 0.16 | 0.04 | 0.23 | 3.19 | 13 | 0.007 |
| 1-4 | 0.14 | 0.11 | 0.08 | 0.21 | 4.88 | 13 | <0.001 |
| 2-3 | 0.00 | 0.15 | -0.08 | 0.09 | 0.10 | 13 | 0.921 |
| 2-4 | 0.01 | 0.12 | -0.06 | 0.08 | 0.38 | 13 | 0.712 |
| 3-4 | 0.01 | 0.13 | -0.07 | 0.08 | 0.24 | 13 | 0.813 |

**Figure 4.**
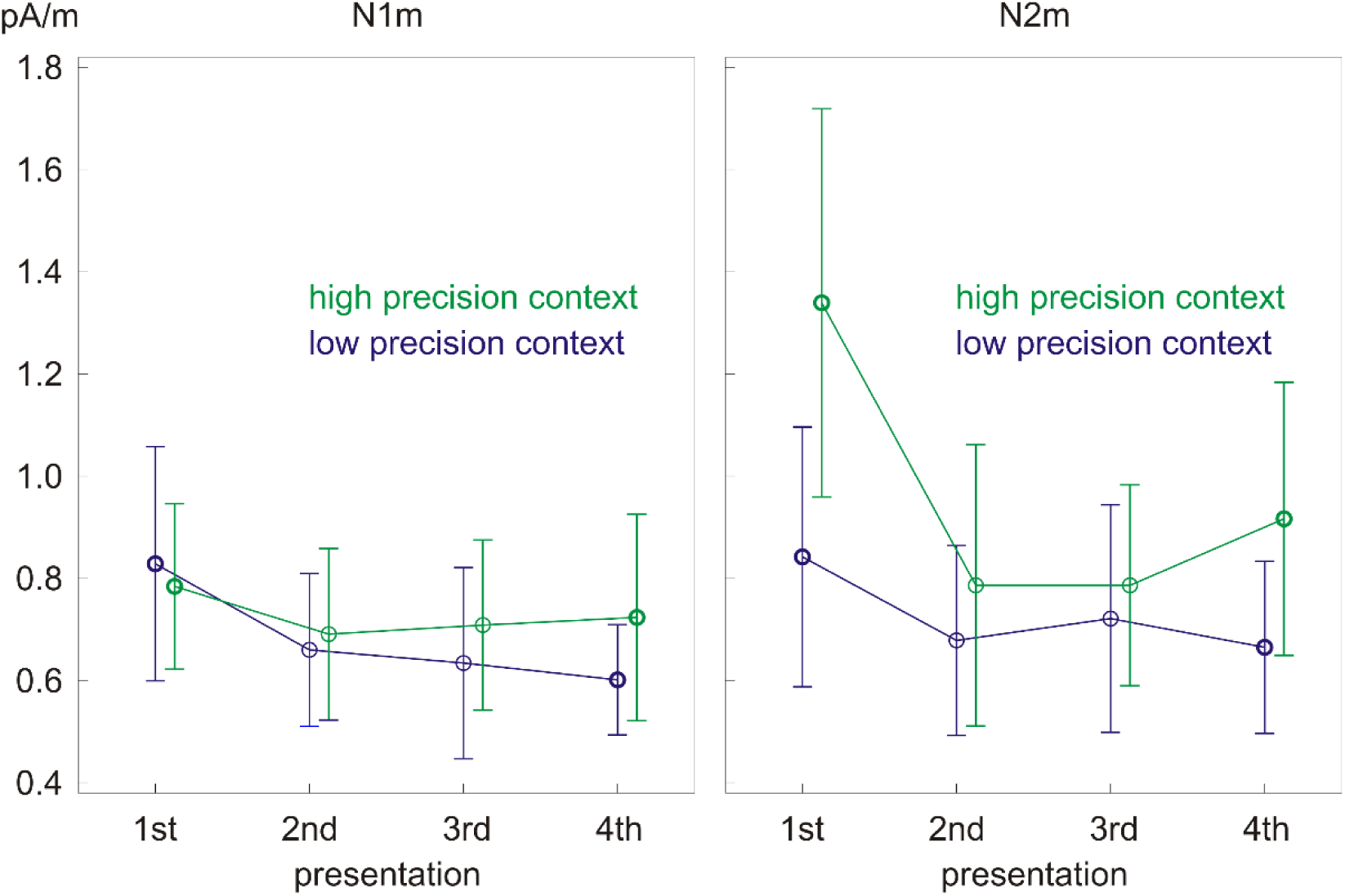
Significant main effect of repetition on N1m source power (left) and precision x repetition interaction on N2m source power (right). Error bars depict one standard deviation of the mean.

The ANOVA on the N2m source power revealed a precision x repetition interaction (F(3,39) = 8.12, p < 0.01, partial eta squared = 0.38) (**Figure 4** right) as well as main effects of precision (F(1,13) = 15.06, p < 0.01, partial eta squared = 0.54) and repetition (F(3,39) = 9.87, p < 0.01, partial eta squared = 0.43, epsilon = 0.48). Post hoc paired samples t-tests looking into the precision x repetition interaction showed that tones in the low precision context had smaller source power than tones in the high precision context upon the 1st presentation (t(13) = -4.24, p < 0.01) and the 4th presentation (t(13) = -3.06, p < 0.01). Moreover, repetition did not modulate the source power in the low precision context but did so in the high precision context (**Table 3**). In the high precision context, the source power decreased from the 1st presentation to the 2nd, 3rd, and 4th presentations (t(13) = 3.75, p < 0.01; t(13) = 4.36, p < 0.01; t(13) = 3.94, p < 0.01). Most interestingly, the source power started to increase again in the 4th presentation as indicated by the difference in source power between the 3rd and 4th presentations (t(13) = -2.26, p < 0.05).

**Table 3.**
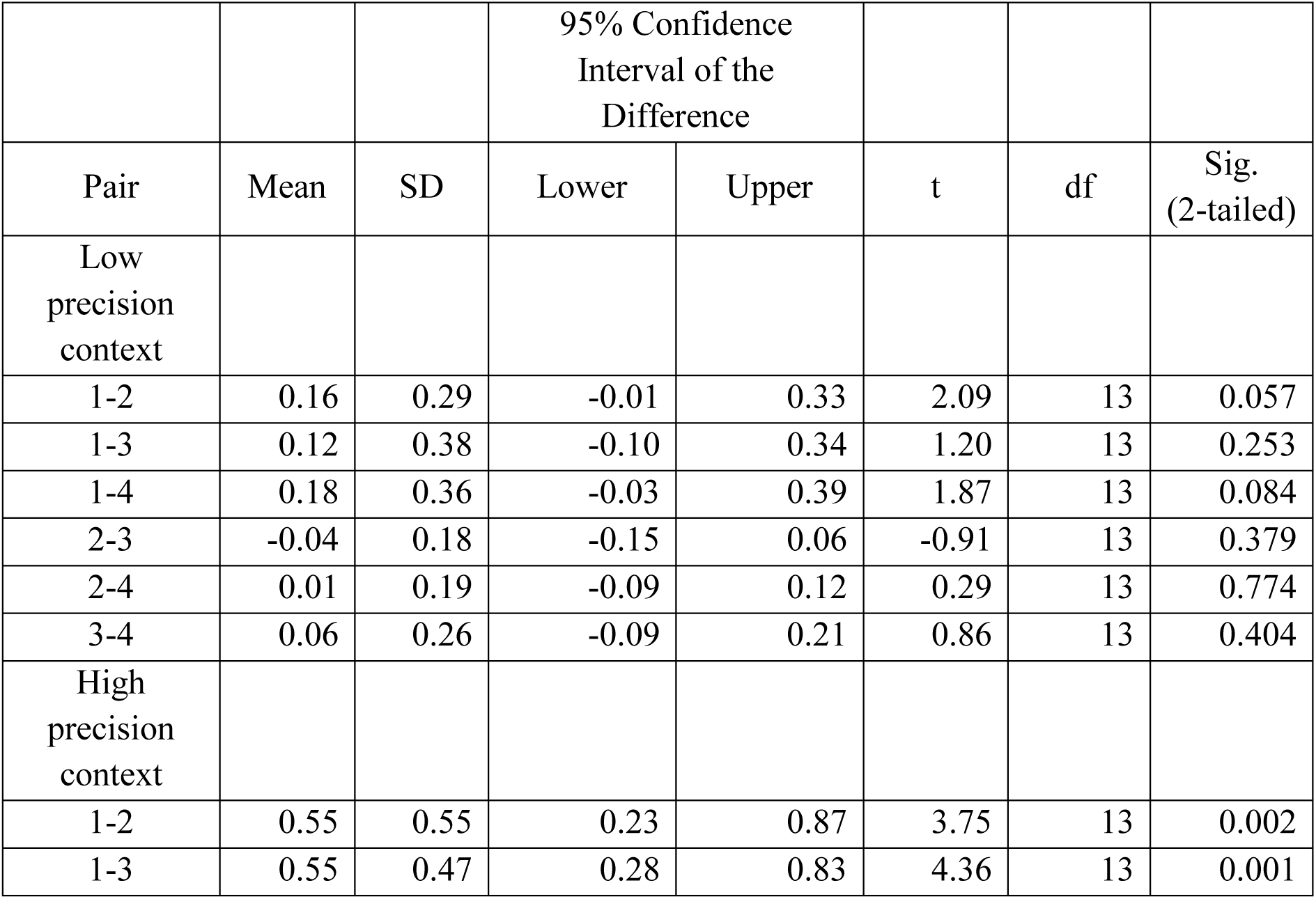
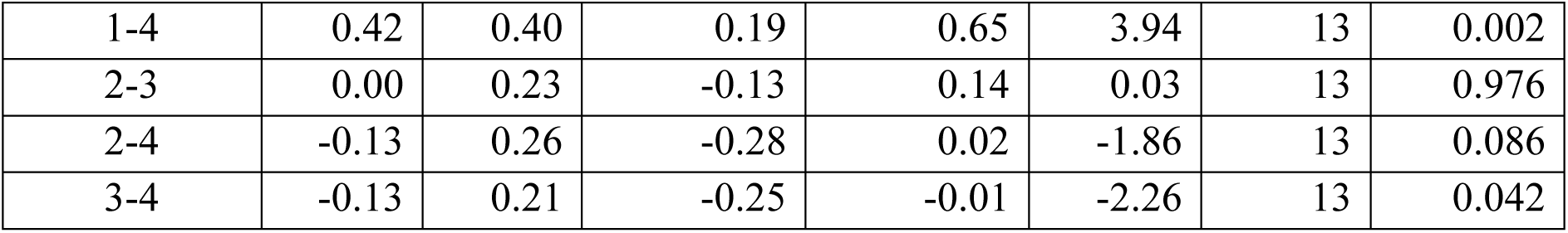
Results of post hoc paired samples t-tests looking into the precision x repetition interaction on N2m source power.

|  |  |  | 95% Confidence Interval of the Difference |  |  |  |  |
| --- | --- | --- | --- | --- | --- | --- | --- |
| Pair | Mean | SD | Lower | Upper | t | df | Sig. (2-tailed) |
| Low precision context |  |  |  |  |  |  |  |
| 1-2 | 0.16 | 0.29 | -0.01 | 0.33 | 2.09 | 13 | 0.057 |
| 1-3 | 0.12 | 0.38 | -0.10 | 0.34 | 1.20 | 13 | 0.253 |
| 1-4 | 0.18 | 0.36 | -0.03 | 0.39 | 1.87 | 13 | 0.084 |
| 2-3 | -0.04 | 0.18 | -0.15 | 0.06 | -0.91 | 13 | 0.379 |
| 2-4 | 0.01 | 0.19 | -0.09 | 0.12 | 0.29 | 13 | 0.774 |
| 3-4 | 0.06 | 0.26 | -0.09 | 0.21 | 0.86 | 13 | 0.404 |
| High precision context |  |  |  |  |  |  |  |
| 1-2 | 0.55 | 0.55 | 0.23 | 0.87 | 3.75 | 13 | 0.002 |
| 1-3 | 0.55 | 0.47 | 0.28 | 0.83 | 4.36 | 13 | 0.001 |
| 1-4 | 0.42 | 0.40 | 0.19 | 0.65 | 3.94 | 13 | 0.002 |
| 2-3 | 0.00 | 0.23 | -0.13 | 0.14 | 0.03 | 13 | 0.976 |
| 2-4 | -0.13 | 0.26 | -0.28 | 0.02 | -1.86 | 13 | 0.086 |
| 3-4 | -0.13 | 0.21 | -0.25 | -0.01 | -2.26 | 13 | 0.042 |

## 1.4. Discussions

The current research used MEG to examine whether prior precision modulates the cortical dynamics of the making of perceptual inferences. We presented participants with cycles of repeated tone quartets which consisted of three prime tones and one probe tone, where the repetition of probe tone can reveal the development of perceptual inferences in low and high precision contexts depending on its position within the cycle. We found that the two conditions modulate the N1m source power in similar manner. However, there was a significant precision x repetition interaction on the N2m source power. While repeated probe tones in low precision context did not exhibit any modulatory effect, repeated probe tones in high precision context were associated with a suppression and a rebound of the N2m source power. The results confirm the necessity to dissociate the processing of non-predicted stimuli of different prior precision (Friston, 2005, 2009; Feldman & Friston, 2010; Arnal & Giraud, 2012; Schröger et al., 2015; Hsu et al., 2015, 2018). Moreover, it is likely that the minimisation of prediction error in low and high precision contexts involves distinct mechanisms.

In electrophysiology literature, N1/N1m is known to reflect multiple processes of signalling unspecific changes in the auditory environment (Näätänen & Picton, 1987; Crowley & Colrain, 2004). Mounting evidence of the N1/N1m predictability effect further supports that it indicates the operation of an internal predictive mechanism, as predicted stimuli were associated with robust N1/N1m suppression (Schafer & Marcus, 1973; Schafer et al., 1981; Lange, 2009; Todorovic et al., 2011; Todorovic & de Lange, 2012; SanMiguel et al., 2013; Timm et al., 2013; Hsu et al., 2014a, 2014b, 2016). Our findings extend prior research by showing that such predictability effects resemble each other in low and high precision contexts, where prediction error is weighted differently. This suggests that changes in N1m source power cannot distinguish between the minimisation of prediction error due to prediction formation (as in low precision context) and prediction alteration (as in high precision context). Instead, N1m seems to reflect the overall reduction in prediction error.

According to the predictive coding model of perception, prediction error can be adjusted depending on the precision of the sensory input (Friston, 2005, 2009; Feldman & Friston, 2010). Prediction error is weighted less in low than high precision contexts (Schröger et al., 2015), leading to smaller N1 responses to target tones following random than regular tone sets in EEG (Hsu et al., 2015). We speculate that the difference between low and high precision contexts might be less conspicuous here so that we did not obtain a main effect of precision on the N1m source power in MEG. In particular, in our previous experiment (Hsu et al., 2015), the difference between the two conditions depended on the regularity of their prime tones. That is, stimuli were preceded by either random or rising tones. However, in the current research, the difference between the two conditions depended on their position within the cycle of stimulus presentation. That is, stimuli were presented in either the beginning or the middle of each cycle. Such procedural differences between investigations might influence how much the two conditions differ in prior precision. This possibility should be systematically tested in future research.

Nevertheless, the differentiation between prediction error processing in low and high precision contexts was evident on the N2m source power. Specifically, tones in low precision context triggered smaller source power than tones in high precision context upon the 1st presentation and the 4th presentation. More importantly, stimulus repetition triggered different response pattern in low and high precision contexts. While repeated tones in low precision context did not exhibit any modulatory effect, repeated tones in high precision context were associated with a suppression and a rebound of the N2m source power. Previous neurocomputational modelling already proposes that stimulus repetition results in perceptual learning which increases the precision of prediction error (Garrido et al., 2009; see Auksztulewicz & Friston, 2016 for a review). Our result pattern further suggests that the increased precision of prediction error can manifest differently at the cortical level depending on its initial precision status.

Specifically, novel probe tones presented in the beginning of each cycle are associated with lower prior precision, as listeners had a general expectation that a probe tone would appear but had little if any idea concerning its frequency. The repetition of these stimuli increases the precision of prediction error, which in turn minimises the cortical responses encoding prediction error. It is possible that this process takes place automatically in the auditory cortices near planum temporale, hence modulating the N1m but not the N2m.

On the other hand, novel probe tones presented in the middle of each cycle are associated with higher prior precision, as listeners already formed predictions on its frequency over previous stimulation within the cycle. These stimuli resemble more the MMN, which is interpreted as a failure to inhibit prediction error due to deviation from a learned regularity (Friston, 2005; Garrido et al., 2007). The repetition of these stimuli also increases the precision of prediction error. However, the information might be propagated into the next hierarchical level, hence modulating the N2m as well. The N2m suppression (from the 1st presentation) is in line with the finding that the MMN vanishes with few stimulus repetitions (Garrido et al., 2009), suggesting that the brain can efficiently adjust a perceptual model. Meanwhile, the N2m rebound (toward the 4th presentation) resonates with the finding of enhanced sustained field at around 200 ms to stimulus repetition (Näätänen & Rinne, 2002; Bendixen et al., 2007; Ylinen & Huotilainen, 2007; Recasens et al., 2015). It supports the notion that factors other than mere probability should be considered in order to account for the way perceptual model modification is implemented in the brain (Hsu et al., 2016). For example, there might be a gradual decrease in the bandwidth of the prediction tuning curve in the high precision context (cf. sharpening model for repetition effect; see Grill-Spector et al., 2006 for a review). The sparser representation of prediction can paradoxically elicit an increase in prediction error. Alternatively, there might be a build-up of representations in prediction alteration. This can introduce a learning function of escalating sensitisation which upweights neuronal responses (Karhu et al., 1997; Barascud et al., 2016; Southwell et al., 2017). Finally, there might be heightened expectation for the onset of a novel pair of tone quartet after the repetition increases in the current pair of tone quartet. Such preparation for change can also account for the suppression-rebound pattern in the high precision context.

Although measures were taken to prevent participants from learning high-order regularities in the current research, it cannot be excluded that participants might become aware of the stimulus structure (i.e., the probe tones would change after 4 to 6 repetitions). However, if this happened, participants would expect for changes of probe tones in both the low and high precision contexts. Therefore, it cannot account for the difference between conditions reported here. It is also unlikely that the dissociation of probe tones in low and high precision context was due to how much the probe tones differ from their preceding tones (i.e., the three prime tones) in terms of frequency. It is because the frequency of these tones was determined by random sampling. The allocation of these tones to low/high precision context was dependent on their position within a cycle (i.e., whether they were presented in the beginning/middle of a cycle) rather than their frequency.

The dissociation of probe tones in low and high precision context is closely related to the mixed results in the literature of repetition-related effects as previously suggested in Hsu et al. (2015). Although repetition-related effects are commonly explained in the language of the Bayesian models of predictive coding (Summerfield et al., 2008), there is a puzzling juxtaposition of repetition enhancement and repetition suppression across functional magnetic resonance imaging (fMRI) research. While the repetition of unfamiliar stimuli was associated with enhanced neuronal responses, the repetition of familiar stimuli was associated with suppressed neuronal responses (Henson et al., 2000; Fiebach et al., 2005; Gruber & Müller, 2005; Gagnepain et al., 2008; Soldan et al., 2008; Subramaniam et al., 2012; Müller et al., 2013). It was proposed that stimuli of different familiarity differ in whether there is a pre-existing representation (Turk-Browne et al., 2008), which can be understood as the top-down activation of predictions (Cheung & Bar, 2012; Grotheer & Kovács, 2014). The repetition of unfamiliar stimuli (initially associated with no pre-existing representation) would resemble the development of perceptual inferences in low precision context. The repetition of familiar stimuli (initially associated with certain pre-existing representation) would resemble the development of perceptual inferences in high precision context. Interestingly, the dissociation of repetition-related effects in the current research did not manifest as repetition enhancement and repetition suppression. Rather, it was expressed as repetition suppression on N1m followed by a lack of modulatory effect on N2m in low precision context versus repetition suppression on N1m followed by a U-shaped profile on N2m in high precision context. Future research is needed to develop theories that relate haemodynamic responses to electrophysiological data.

## Acknowledgements

This work was supported by Taiwan Ministry of Science and Technology (grant number 105-2410-H-003-145-MY3) to YFH and Department of Psychology at the University of Jyväskylä and Academy of Finland (profiling action “Multilete” #292466) to JAH. We thank Mr. Lauri Kantola for assistance with MEG data collection.

